# Dietary lysozyme supplement alters serum biochemical makers and milk metabolite profile of sows via gut microbiota

**DOI:** 10.1101/444778

**Authors:** Jian Zhou, Xia Xiong, Lijun Zou, Jia Yin, Kexing Wang, Yirui Shao, Yulong Yin

## Abstract

Lysozyme is an important antimicrobial agent with promising future in replacing antibiotics in livestok production. The aim of current study was to determine variations in sow’s gut microbiota, serum immunity and breast milk metabolite profile mediated by lysozyme supplementation.Thirty-six pregnant sows were assigned to a control group without supplementation and two treatments with 0.5 g/kg and 1.0 g/kg lysozyme provided in formula feed for 21days. Microbiota analysis based on 16s RNA high-throughput sequencing and untargeted liquid chromatography tandem mass spectrometry were applied and combined in analysis. Serum biochemical indicators and immunoglobulins were also determined. Sows received 1.0kg/t lyszoyme treatment shown significant redution in microbial diversity. Spirochaetes, Euryarchaeota and Actinobacteria significantly increased while Firmicutes showed a remarkable reduction in 1.0kg/t treated group compared with control. Pyrimidine metabolism,Purine metabolism and Amino acid related enzymes were significantly upregulated in 1.0kg/t lysozyme treated group. The richness of gram-positive bacteria were significantly down-regulated by lysozyme treatments. Serum aspartate transaminase (AST) activity was significantly un-regulated. Serum IgM levels were significantly higher in the 1.0 kg/t group compared with control, while IgA levels was significantly lower in 1.0kg/t group. Over thirty metabolites from sow’s breast milk including L-Glutamine,creatine and L-Arginine were sigficantly altered by lysozyme treatment. There existed crucial correlations among gut microbiota, serum immunity and breast milk metabolites where lactobacillus and prevotella may play a key role in lysozyme mediated host-microbial interactions. Overall, lysozyme supplementation could effectively improve the composition, metabolic functions and phenotypes of sow’s gut microbiota and it also benefit sows with better immune status and breast milk composition.

**Importance:** Enteric infections caused by pathogens have a significant negative effect on neonatal survival and animal health in swine production. The application of antibiotics in feeds at subtherapeutic levels could improve performance and overall health and is used extensively throughout the industry. However, abuse of antibiotics is contributing to the high level of drug resistance in microbial communities and rising concerns regarding human health. Here, we revealed that lysozyme supplementation could effectively improve the composition, metabolic functions and phenotypes of sow’s gut microbiota and it also benefit sows with better immune status and breast milk composition. These findings confirmed that lysozyme could be a suitable alternative to antibiotics in swine production.

## 1. Introduction

Enteric infections caused by pathogens, like Enterotoxigenic *Escherichia coli* (ETEC), have a significant negative effect on neonatal survival and animal health in swine production [1, 2, 3]. Animal infants infected by pathogenic bacteria often suffer from persistent diarrhea and serious inflammation [3, 4]. Prolonged inflammation of the intestinal tract leads to substantial destruction of the intestinal epithelia, resulting in malnutrition and impairing the early growth of infants [5, 6, 7]. Application of antibiotics in formula feed is well established method and can improve growth rates of piglets [8]. However, abuse of antibiotics is contributing to the high level of drug resistance in microbial communities and rising concerns regarding human health [1, 6, 9, 10]. An alternative to antibiotics is lysozyme, an enzyme and natural broad-spectrum fungicide commonly found in tears, saliva, and milk, and that is a vitally important immune system activator under physiological conditions [11, 12, 13]. During bacterial infection of the intestine, mammalian Paneth cells are also able to secrete lysozyme via secretory autophagy to maintain intestinal homeostasis [14].

Breast milk contains lysozyme (< 0.065 μg/mL), along with lactoferrin and secretory IgA (SIgA), which greatly aid the establishment of beneficial gut microbiota in newborns [11, 15]. Lysozyme functions by cleaving the β-1,4-glycosidic bond between N-acetylmuramic acid and N-acetylglucosamine residues of the bacterial peptidoglycan, causing a loss of cellular membrane integrity and cell lysis [1, 10]. Lysozyme was reported to be more effective against gram-positive bacteria [16, 17, 18], and it may also indirectly affect several gram-negative bacterial species [12, 19]. Previous studies revealed that lysozyme promotes the beneficial microbe community and reduces detrimental microbes within gut microbial communities [11, 20]. Lysozyme could be effective against a wide range of gastrointestinal pathogens, such as *Listeria monocytogenes*, *Clostridium perfringens*, *Candida* spp., and *Helicobacter pylori* in vitro [7, 21]. Bacterial sensitivity to lysozyme is due to the activation of innate components of the immune system, such as increased neutrophil activation during inflammation [3, 22]. It has been reported that lysozyme may possess an anti-inflammatory effect via inhibiting JNK phosphorylation [23]. Furthermore, lysozyme is capable of enhancing intestinal SIgA secretion, cause macrophage activation, and promote rapid clearance of bacterial pathogens [2, 13, 22].

The most important characteristics for any antimicrobial feed additives are weight gain and feed efficiency. Recent studies reported that lysozyme sourced from chicken eggs showed significant advances in improving growth performance, intestinal morphology, gut microbiota composition, and immunity of piglets [1, 2, 15, 24, 25]. For instance, weaned piglets received a hen-egg white lysozyme treatment shown better intestinal growth and development, as well as decreased ETEC counts on the intestinal mucosa and serum proinflammatory cytokines [26]. Moreover, lysozyme produced by transgenic animals and structurally modified lysozyme was shown to possess significant antimicrobial properties against pathogens like ETEC in piglets [12, 21, 26, 27, 28, 29]. Piglets that consumed lysozyme-transgenic goats’ milk (containing human lysozyme at 67% of the concentration in human breast milk) showed better intestinal morphology and fewer total coliform counts [21]. These recent studies confirmed the promising future of lysozyme in replacing antibiotics.

Gut microbiota plays multiple roles in animal growth and health, including energy extraction from the diet, gut barrier function and immune system maturation, and growth performance [30, 31]. However, none of these studies above provided a systematic overview of lysozyme mediated variations in gut microbiota and its potential interactions with immune systems. Moreover, increasing evidence suggests that maternal diet during pregnancy modifies an offspring’s microbiota composition and intestinal development in a long-term manner [32, 33]. Nutritional intervention on sows with additives results in greater neonatal survival and infant health [15]. Given this, in the present study, 36 pregnant sows were assigned to a control group without supplementation and two treatments with 0.5 g/kg and 1.0 g/kg lysozyme provided in formula feed. After the 21-day supplementation, the effects of lysozyme on sow’s gut microbiota, breast milk metabolite profile, and serum biochemical indices were systematically investigated and associations among them mediated by lysozyme treatment were also revealed for the first time.

## Materials and methods

### Animals and ethics statement

All procedures involving animals were carried out in accordance with guidelines for animal studies issued by the Animal Care and Use Committee of the Institute of Subtropical Agriculture, Chinese Academy of Sciences [34]. Modified hen-egg white lysozyme additives were obtained from Shanghai E. K. M Biotechnology Co. Ltd., Shanghai, China. A total of 36 multiparous hybrid pregnant sows (Landrace×Yorkshire) with an average parity of 4.67±1.50 were selected to this study and then randomly assigned to three groups (n = 12 per treatment), including a control group (CN) without supplementation and two treatments with 0.5 g/kg (LA) and 1.0 g/kg (LB) lysozyme provided in formula feed. The current study started 24 days before the expected date of confinement. Sows did not have diseases like diarrhea and had never received antibiotics before the study. Lysozyme supplementation continued for 21 days till prenatal fasting. Breast milk from all investigated sows were collected on the due date.

### Sample collection and processing

After the 21-day supplementation, fresh feces from each individual were collected on the same day using 5 mL sterile centrifugal tubes, immediately frozen in liquid nitrogen, and stored at −80°C until DNA extraction. After the 21-day supplementation, the sows were restrained for blood sampling and approximately 5 mL blood was collected in a vacuum tube from the sow’s auricular vein and directly centrifuged at 1500 × *g* for 15 min. The supernatant of each sample was then divided into subsamples and stored at −20°C until analysis. On parturition day, breast milk from each individual was collected with 5 mL sterile centrifugal tubes and immediately frozen at −20°C until further analysis.

### Microbiota analysis based on 16S RNA high-throughput sequencing

Eight fecal samples from sows in each group (n =8 per treatment) were randomly chosen for microbiota analysis and total bacterial DNA was extracted from approximately 0.25 g of feces using a QIAamp DNA Stool Mini Kit (Qiagen, Hilden, Germany) according to the manufacturer’s instructions. The diversity and composition of the bacterial community were determined by high-throughput sequencing of the microbial 16S rRNA genes. The V4 hypervariable region of the 16S rRNA genes and was PCR amplified using 515F: 5′-GTGCCAGCMGCCGCGGTAA-3′ and 806R: 5′-GGACTACHVGGGTWTCTAAT-3′ primers, Illumina adaptors, and molecular barcodes. Paired-end sequencing was performed on the Illumina HiSeq 2500 platform (Novogene, Beijing, China). Raw 16S data sequences were obtained before being screened and assembled using the QIIME (v1.9.0) [35] and FLASH software packages. UPARSE (v7.0.1001) [36] was used to analyze the high-quality sequences and determine OTUs. Subsequently, high-quality sequences were aligned against the SILVA reference database (https://www.arb-silva.de/) and clustered into OTUs at a 97% similarity level using the UCLUST algorithm (https://drive5.com/usearch/manual/uclust_algo.html). Each OTU was assigned to a taxonomic level with the Ribosomal Database Project Classifier program v2.20 (https://rdp.cme.msu.edu/). The assembled HiSeq sequences obtained in the present study were submitted to the NCBI’s Sequence Read Archive (SRA, No. PRJNA415259) for open access.

### Metagenomic prediction and metabolic phenotype analysis

Functional metagenomes of all samples were predicted using PICRUSt v1.1.3 [37]. OTUs were determined according to the instructions provided in the Genome Prediction Tutorial for PICRUSt. Metagenomes (http://picrust.github.io/picrust/) were predicted from the copy number normalized 16S rRNA data in PICRUSt using the predict metagenomes.py script against the functional database of KEGG Orthology. Functional categories at different levels were computed with the script categorize by function.py. Functional differences within groups were explored using LEfSe and specific analysis was performed through the Galaxy server [38]. Output files from the PICRUSt analysis were collected and analyzed by R software (v3.5.1) for further statistical interrogation and graphical depictions of all predicted functional datasets. BugBase (https://bugbase.cs.umn.edu/) was employed to predict organism-level microbiome phenotypes using 16S RNA datasets and mapping file according to the tutorial [39].

### Serum biochemical indices and immunoglobulins analysis

Eight serum samples from each group (n = 8 per treatment) was randomly chosen for further analysis of biochemical and immune indices. Serum parameters investigated in the present study included: total protein (TP), blood urea nitrogen (BUN), creatinine (CREA), cholesterol (CHO), triglycerides (TG), high-and low-density lipoprotein cholesterol (HDL-C, LDL-C), glucose (GLU), albumin (ALB), globulin (GLO), AST, and ALT. All parameters were measured using the TBA-120FR biochemistry analyzer provided by the Biochemical Analysis Center of Hunan Normal University Hospital. IgG, IgM, and IgA were measured with enzyme-linked immunosorbent assay kits (Cusabio Biotech Co., Hubei, China) according to the manufacturer’s instructions and previous research [40].

### Untargeted metabolomic analysis

Breast milk from sows with nearby delivery time (Within 48 hours, n = 6 per treatment) were chosen for untargeted metabolomic analysis based on the Liquid Chromatography Tandem Mass Spectrometry (LC-MS/MS) platform. Milk samples were slowly thawed at 4°C and then 100 μl of each sample was added to 400 μl pre-cooled methanol/acetonitrile (1:1, v/v), vortex mixed, stood at −20°C for 60 min, centrifuged at 14,000 *g* for 20 min at 4°C, take the supernatant and vacuum dried. For mass spectrometry, 100 μl of acetonitrile aqueous solution (acetonitrile:water = 1:1, v/v) was reconstituted, vortexed, centrifuged at 14,000 g for 5 min at 4°C, and the supernatant was taken for further analysis on LC-MS/MS platform (Bioprofile Co. Ltd, Shanghai, China). Each sample was tested by positive ion and negative ion mode using electrospray ionization (ESI). Samples were separated by Ultra Performance Liquid Chromatography (UPLC) and analyzed by mass spectrometry using a Triple-TOF 5600 mass spectrometer (AB SCIEX). The raw data was converted to .mzXML format by ProteoWizard [41], and then the XCMS program [42] was used for peak alignment, retention time correction, and peak area extraction. Metabolomic data were analyzed with MetaboAnalyst v4.0 [43] online version (http://www.metaboanalyst.ca/faces/home.xhtml). Key metabolites were filtered by VIP scores and rules used in our previous research [44].

### Statistical Analysis

All statistical analyses were performed using SPSS 25.0 software (SPSS Inc., Chicago, IL). Alpha and beta diversity were analyzed with QIIME (v1.7.0) and displayed with R software (v3.5.1) and details can be found in the legends of the corresponding figures and tables. The differences among groups were compared using one-way ANOVA and Tukey-Kramer multiple comparison tests. *P* values < 0.05 were used to indicate statistical significance, whereas 0.05 ≤ *P* values < 0.1 were considered to be trending toward significance.

## Results

### Dietary lysozyme supplementation altered the diversity and composition of sow’s gut microbiota

Microbial diversity, evidenced by the Shannon index, showed a significant reduction in the 1.0 kg/t group compared with the lysozyme-treated group (p = 0.0014, Figure 1A) and no remarkable differences were found in indicators of microbial richness (ACE and Chao1, Sup Table 1). The principal coordinate analysis (PCoA) based on Bray-Curtis dissimilarity revealed that microbiota showed obvious segregation from the control group to lysozyme-treated groups (Figure 1B). In addition, non-metric multidimensional scaling (NMDS) plots of β-diversity weighted unifrac (Figure 1C) also confirmed the differences between control and lysozyme-treated groups (all P < 0.05 by Anosim analysis and multi-response permutation procedure (MRPP)). Furthermore, an unweighted pair-group method with arithmetic mean (UPGMA) analysis based on weighted unifrac distances were applied and the phylogeny showed the relationships of all observed samples. The phylogeny revealed that *Firmicutes*, *Bacteroidetes*, *Proteobacteria,* and *Fibrobacteres* are the dominant bacteria in sow’s gut microbiome (Sup Figure1).

**Figure.1.**
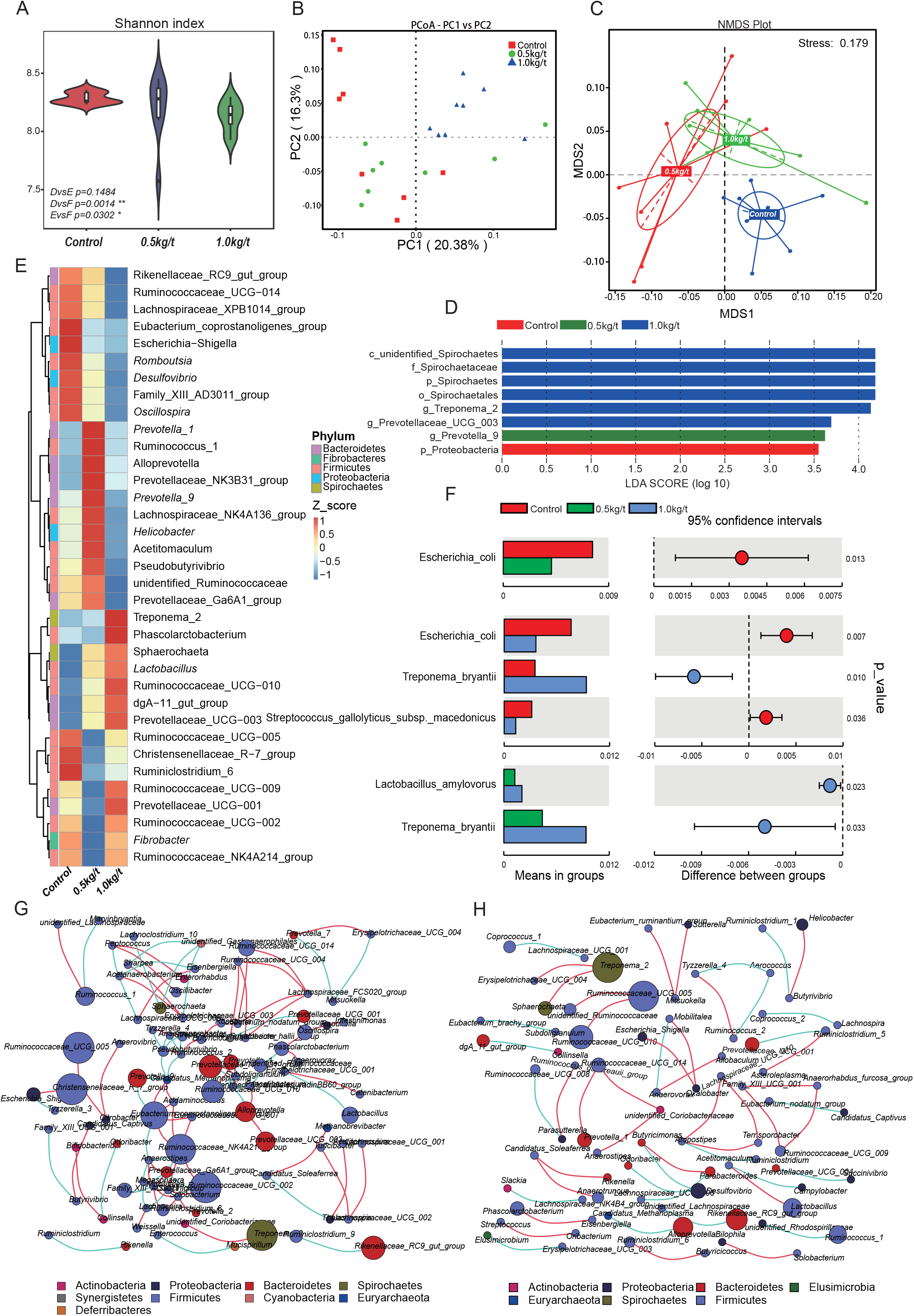
Effects of different lysozyme levels on sow’s gut microbiota. **A.** The microbial alpha diversity (as accessed by Shannon index) based on whole OTU table. The distribution and density of samples are dispalyed in violin plot. Boxes represent the interquartile ranges, the inside black polts represent the median, and circles are outliers. P values are from Wilcoxon rank sum test. **B-C.** Principal coordinate analysis (PCoA) and non-metric multidimensional scaling (NMDS) analysis based on the OTU table. Significant P values of Anosim and multi-response permutation procedure (MRPP) between groups emphasize the differences in microbial community structure. **D.** LEfse analysis at different microbial taxonomic levels (linear discriminant analysis (LDA score=3.5)). **E.** Heatmap tree shows genera significantly different among groups and their phylogenic relationships. The abundance profiles are expressed by z-scores, and genera were clustered based on Bray Curtis distance in the clustering tree. **F.** T-test bar plot of significantly differed species between groups. **G** and **H**. Spearman’s correlation networks based on genera profile. Control group (**G**) and 1.0kg/t treated group (**H**) showed alterations in microbial relationships.

Further, variations in the microbial composition of all groups were explored. LEfSe analysis of the bacterial community was used to filter the significantly different OTUs among groups and the results showed that there exist dramatic differences in microbial composition between the 1.0 g/kg group and the control group (Figure 1D). *Spirochaetes*, *Euryarchaeota,* and *Actinobacteria* significantly increased while *Firmicutes* showed a remarkable reduction in the 1.0 kg/t treated group compared with the control. *Proteobacteria* also showed lower richness in the 1.0 kg/t group (p = 0.077, Table 1). The heat map (according to the top 35 most different genera) shows the taxonomic distributions among each group (Figure 1E). Specifically, *Escherichia coli* was dramatically reduced in both the 0.5 kg/t and the 1.0 kg/t lysozyme-treated groups. Furthermore, *Lactobacillus amylovorus* showed a significant increase in the 0.5 kg/t group (Figure 1F).

**Table 1.** Changes in major microbial phylum of sows’ gut microbiota after the 21-day lysozyme supplementation^1^

| Taxonomy (%) | Control | 0.5kg/t | 1.0kg/t | *p*-value |
| --- | --- | --- | --- | --- |
| Firmicutes | 63.08<sup>a</sup> | 59.27<sup>ab</sup> | 57.09<sup>b</sup> | 0.033 |
| Bacteroidetes | 26.79 | 31.07 | 30.02 | 0.141 |
| Spirochaetes | 4.42<sup>b</sup> | 4.95<sup>b</sup> | 7.57<sup>a</sup> | 0.011 |
| Proteobacteria | 3.33 | 2.91 | 2.67 | 0.077 |
| Fibrobacteres | 0.76 | 0.44 | 0.73 | 0.761 |
| Euryarchaeota | 0.30<sup>ab</sup> | 0.14<sup>b</sup> | 0.52<sup>a</sup> | 0.010 |
| Actinobacteria | 0.36<sup>ab</sup> | 0.30<sup>b</sup> | 0.43<sup>a</sup> | 0.023 |
| Cyanobacteria | 0.34 | 0.30 | 0.37 | 0.418 |
| Tenericutes | 0.34 | 0.31 | 0.30 | 0.549 |
| Elusimicrobia | 0.05 | 0.06 | 0.08 | 0.279 |
| Others | 0.23 | 0.25 | 0.24 | 0.806 |

<sup>1</sup> Data were analyzed by one-way ANOVA followed by Tukey-Kramer multiple comparison tests.
<sup>a, b</sup> Values within a row with different superscripts differ significantly at $p < 0.05$ .

To further determine the relationships among different microbes in the control and the 1.0 kg/t group, network analysis of gut microbiome was determined by calculating Spearman’s correlation coefficients among all genera. The results revealed that lysozyme supplementation with 1.0 kg/t rebuilt interactions among different genera (Figure 1H). Compared with the control (Figure 1G), gut microbiota shaped by 1.0 kg/t lysozyme treatments had fewer cross-linking, more positive correlations and shorter interactions indicated by lower graph densities [GD] (0.0037 and 0. 0027 in the control and 1.0 kg/t group, respectively), lower average degree [AD] (1.64 and 1.17), lower network diameters [ND] (5 and 4), lower average path lengths [APL] (1.66 and 1.17), higher modularity [MD] (0.87 and 0.94), and higher cluster coefficients [CC] (0.48 and 0.66).

### Lysozyme addition shifted the metabolic functions of sow’s gut microbiota

To investigate further changes in metabolic functions in gut microbiota driven by lysozyme treatment, Picrust was used to predict the metagenome based on 16S RNA sequencing results. At KEGG level 2, Forty functional pathways like amino acid metabolism, carbohydrate metabolism and lipid metabolism showed significant variations among groups (Figure 2A). The principal component analysis (PCA) based of KEGG annotation results revealed that the metabolic functions of sow’s gut microbiota showed obvious segregation from the control group to lysozyme-treated groups (Figure 2B). Furthermore, 80 pathways were found to significantly differ among groups at KEGG level 3, including those associated with cellular processes, environmental factors, genetic information processing, organismic systems, metabolism, and human diseases (Table S2). LEfSe analysis of the KEGG annotation results was used to filter the significantly differed pathways among groups and results showed that there exist dramatic differences in microbial composition between the 1.0 g/kg group and the control group (Figure 1D), which is in line with variations in microbial structure. In the present study, metabolism related pathways at KEGG Level 3 were specifically concerned and filtered. The heat map (according to the most different metabolism related pathways) showed the specific functional pathway distribution among each group (Figure 2D). Moreover, pyrimidine metabolism, purine metabolism and amino acid related enzymes were significantly upregulated in the 1.0 kg/t lysozyme-treated group.

**Figure.2.**
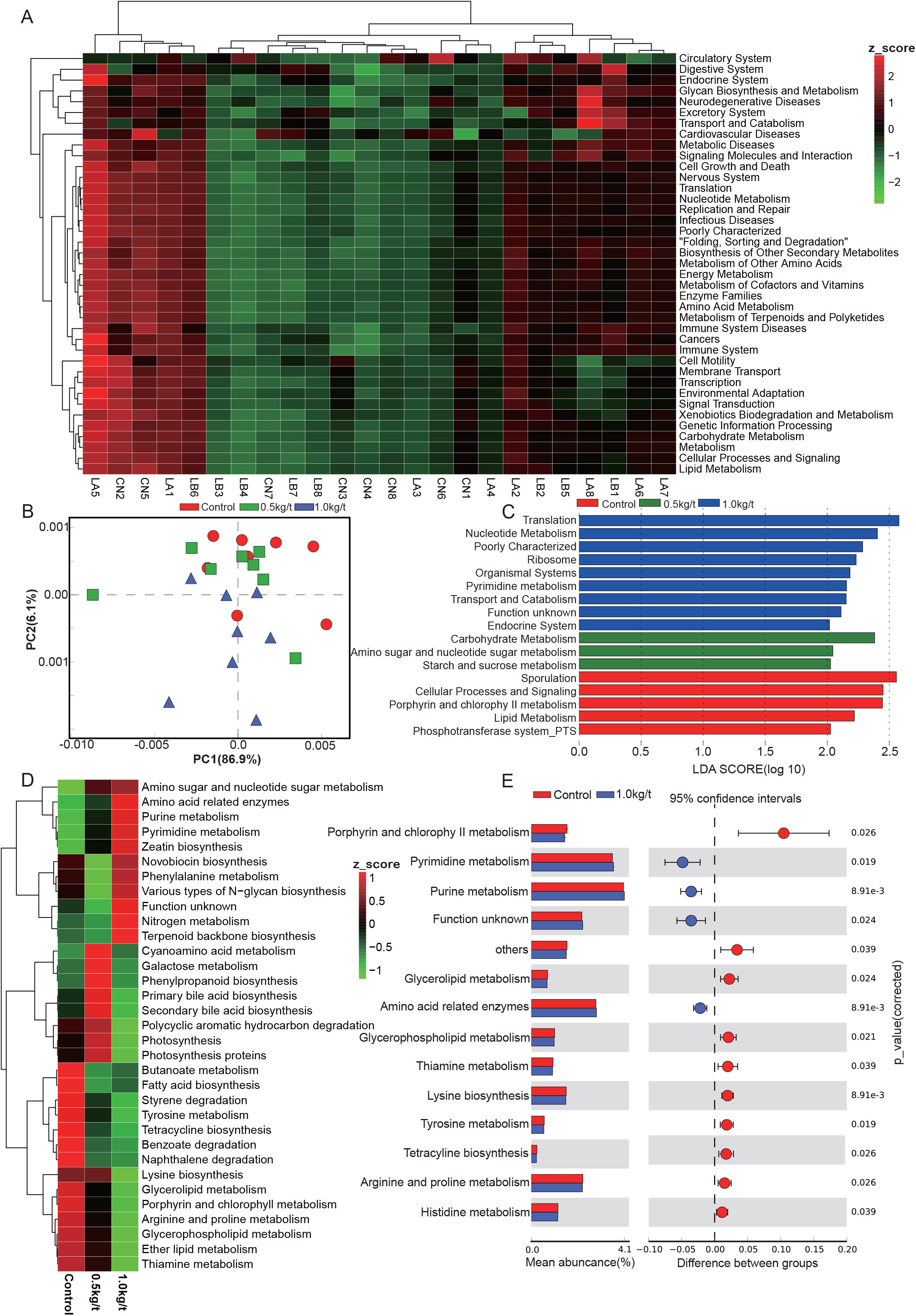
Lysozyme supplementation shifted the metabolic functions of sow’s gut microbiota. **A**. Heatmap tree shows functional profile significantly different among groups at KEGG level 2and their phylogenic relationships. The abundance profiles are expressed by z-scores, and pathways were clustered based on Bray Curtis distance in the clustering tree. **B.** Principal components analysis (PCA) plot of functional profiles upon lysozyme supplementation. **C.** LEfse analysis at different KEGG taxonomic levels (linear discriminant analysis (LDA score=2.0))**D.** Heatmap tree shows metabolism related pathway significantly different among groups at KEGG level 3 and their phylogenic relationships. **E.** T-test bar plot of significantly differed pathways between 1.0kg/t treated groups and control group at KEGG level 3.

### Lysozyme treatment significantly reduced the richness of Gram positive bacteria

To determine the reported impact of lysozyme on gram-positive bacteria, organism-level coverage of functional pathways and biologically interpretable phenotypes were predicted via BugBase, an algorithm that predicts organism-level coverage of functional pathways, as well as biologically interpretable phenotypes, such as oxygen tolerance, gram staining and pathogenic potential, within complex microbiomes using either whole-genome shotgun or marker gene sequencing data. Results demonstrated that the richness of gram-positive bacteria were significantly down-regulated by lysozyme treatments, while gram-negative bacteria showed a significant increase (Figures 3D and 3E). Mobile genetic elements and oxidative stress tolerance of gut microbiota were reduced by increased lysozyme level (Figures 3F and 3I). Otherwise, lysozyme supplementation significantly increased biofilm formation in the lysozyme-treated groups (Figure 3H).

**Figure.3.**
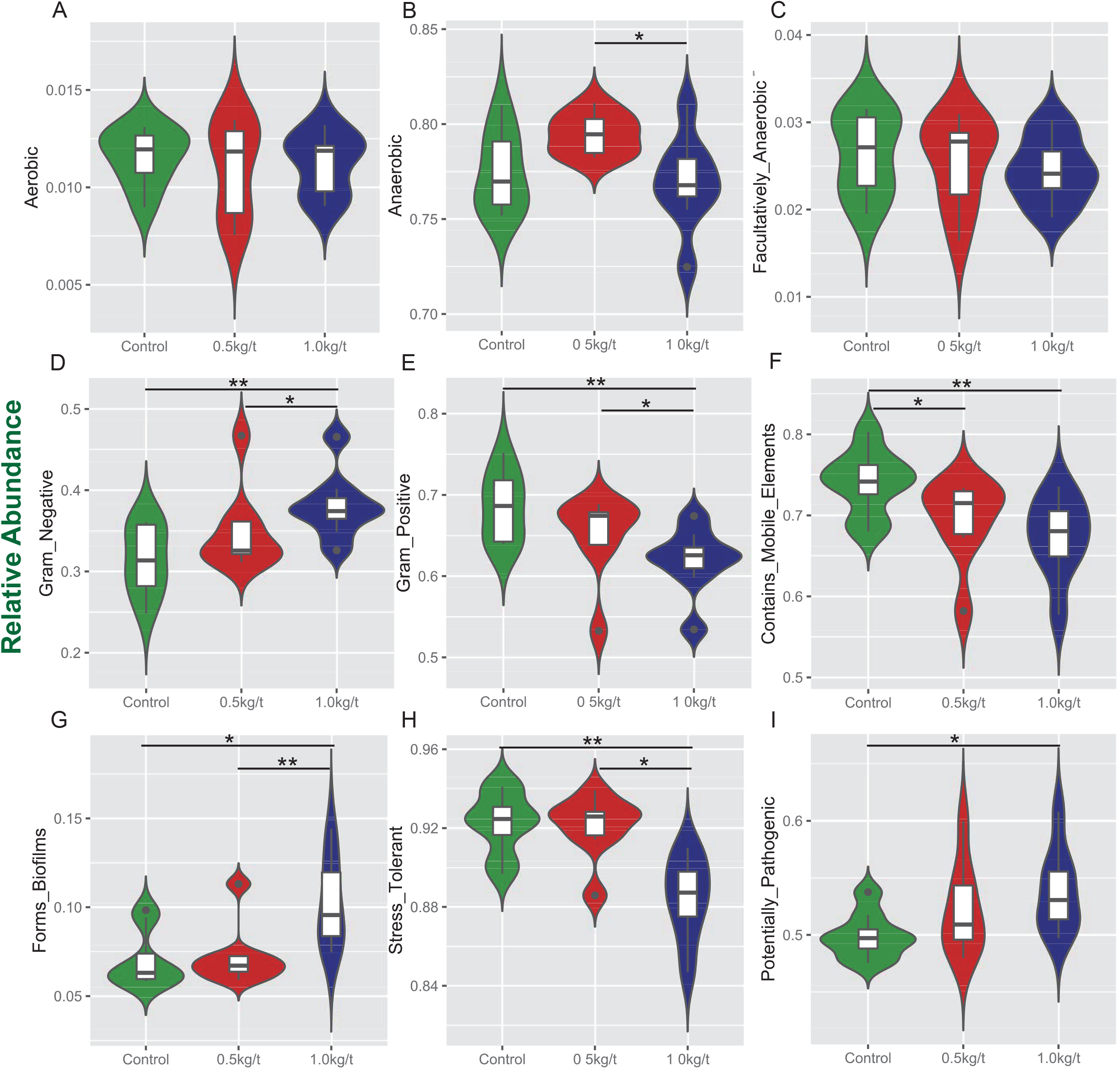
Variations in metabolic phenotypes of sows’ microbiota driven by lysozyme treatment. These results were generated from Bugbase online (https://bugbase.cs.umn.edu/). **A**–**C**. Oxygen utilizing. **D**–**E**. Gram bacterial classification. **F**. Mobile-element containing. **G**. Biofilm forming. **H**. Oxidative stress tolerant. **I**. Potential pathogenic risk. Discrete phenotype relative abundances were compared using pair-wise Mann-Whitney U tests with false discovery rate correction, *P <0.05, **P <0.01.

### Lysozyme mediated changes in serum biochemical markers and their correlations with gut microbiota

To identify the impact of lysozyme on sows’ immunity, serum biochemical indices and immunoglobulins were determined. Results showed that serum aspartate transaminase (AST) levels were significantly upregulated by lysozyme treatment and as a result, AST/ALT (alanine transaminase) ratios rose in the lysozyme-treated group (p = 0.080) (Table 2). Dietary lysozyme supplementation had no significant impact on serum metabolite profiles (e.g., HDL-c, LDL-c, TG, and BUN). For immunoglobulins, serum IgM levels were significantly higher in the 1.0 kg/t group compared with the control, while IgA levels were significantly lower in the 1.0 kg/t group (Figure 4A and C).

**Table 2.**
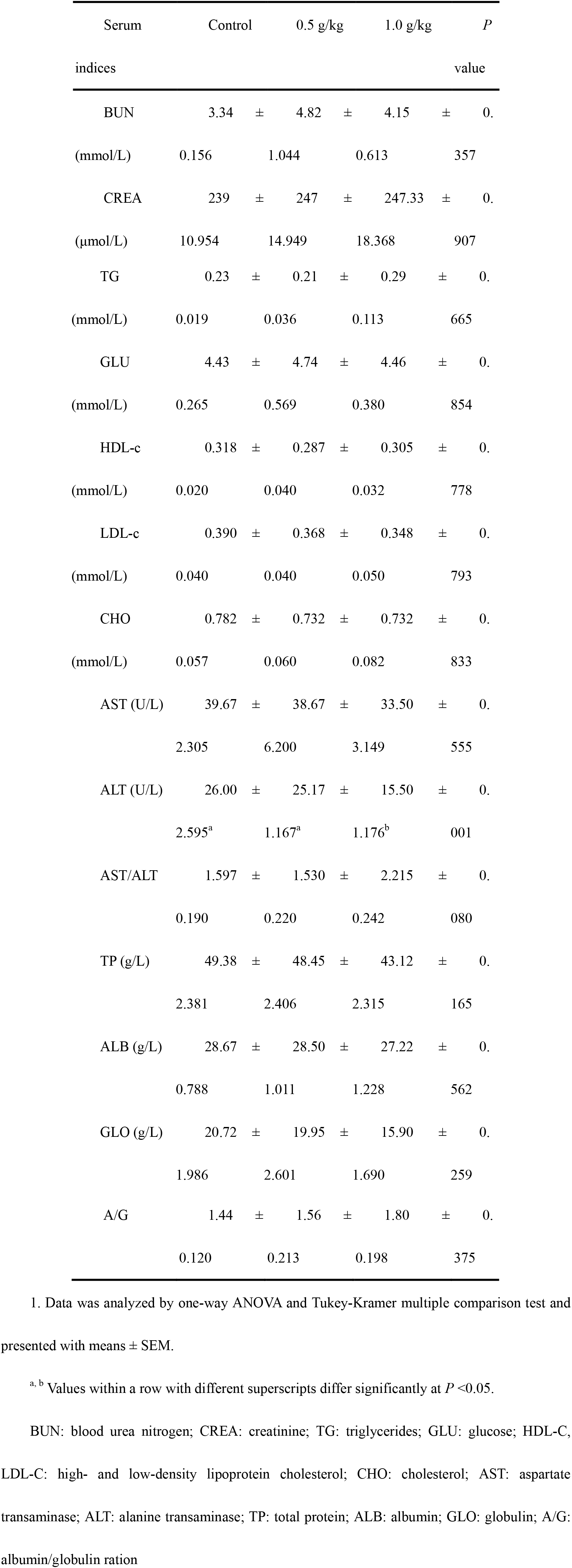
Effects of different lysozyme levels on sows’ serum biochemical indices^1^

**Figure.4.**
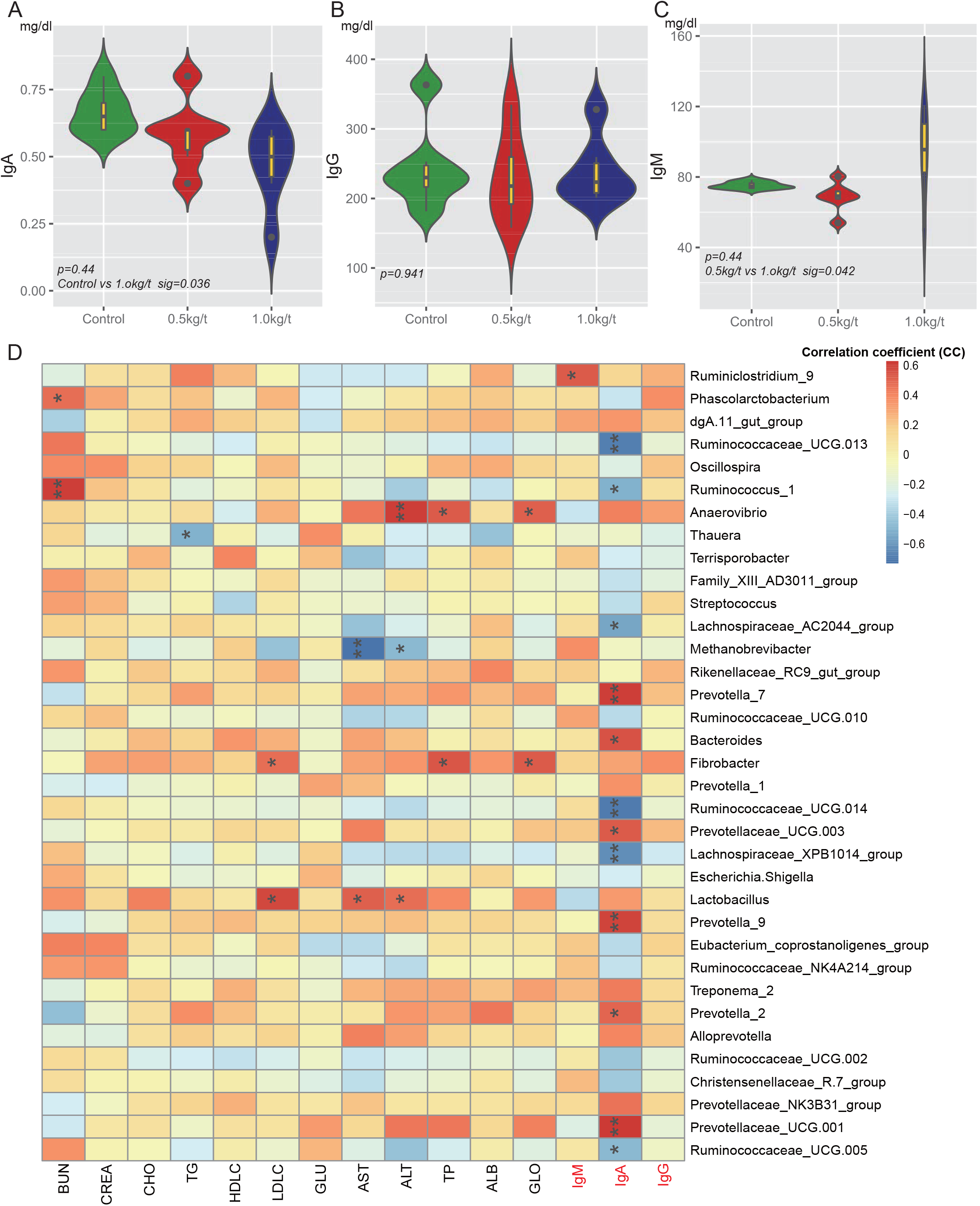
Lysozyme mediated changes in serum Immunoglobulins and correlations with gut microbiota. **A**-**C**. The distribution and density of samples are dispalyed in violin plot. Boxes represent the interquartile ranges, the inside black polts represent the median, and circles are outliers. P values are from one-way ANOVA test. **D.** Heatmap of the Spearman r correlations between the gut microbiota significantly modified serum biochemical makers after 21-day lysozyme supplementation. *P <0.05, **P <0.01 (following the Spearman correlation analysis). BUN: blood urea nitrogen; CREA: creatinine; TG: triglycerides; GLU: glucose; HDL-C, LDL-C: high-and low-density lipoprotein cholesterol; CHO: cholesterol; AST: aspartate transaminase; ALT: alanine transaminase; TP: total protein; ALB: albumin; GLO: globulin; A/G: albumin/globulin ratio.

To further explore the relationship between immunity and the altered gut microbiome driven by lysozyme treatment, Spearman’s correlation coefficients between serum biochemical makers and immunoglobulins and major genera were calculated and visualized with heatmaps. Twelve genera, including *Prevotella*, *Ruminococcaceae UGG*, and *Bacteroides* showed significant correlations with IgA (Figure 4D). *Ruminiclostridium 9* showed a significant positive relation with IgM. *Methanobrevibacter* showed a significantly negative relationship with AST, while *Lactobacillus* showed a significant positive correlation (Figure 4D).

### Variations in sow’s breast milk metabolite profile driven by different lysozyme levels

To evaluate the effects of lysozyme treatment on sow’s breast milk, untargeted Liquid Chromatography Tandem Mass Spectrometry (LC-MS/MS) was applied to assess the metabolite profiles after the 21-day supplementation. Principal component analysis (PCA) of metabolites cannot effectively distinguish the variations among groups (Figure 5A), therefore, Partial Least Squares - Discriminant Analysis (PLS-DA) and Sparse Partial Least Squares - Discriminant Analysis (sPLS-DA) were used (Figure 5B and 5C). Both techniques revealed a distinct partition between lysozyme-treated groups and the control. The heat map (according to the most different metabolites) showed specific metabolite distributions within each group (Figure 5D). Twenty metabolic makers were identified by PLS-DA including Uridine 5′-diphosphate (UDP), UDP-D-glucuronate, acamprosate, and triethanolamine (Figure 5E). In addition, ten metabolites were filtered by sPLS-DA including succinate, L-glutamine, and UDP-D-glucuronate (Sup Figure 2). Given this, metabolites significantly differed among groups were filtered and combined via both PLS-DA and sPLS-DA (Table 3). Moreover, significant differences metabolites between the 1.0 kg/t and the control groups were also filtered via PLS-DA (Sup Figure3) and metabolic makers were obtained (Sup Table. 4).

**Figure.5.**
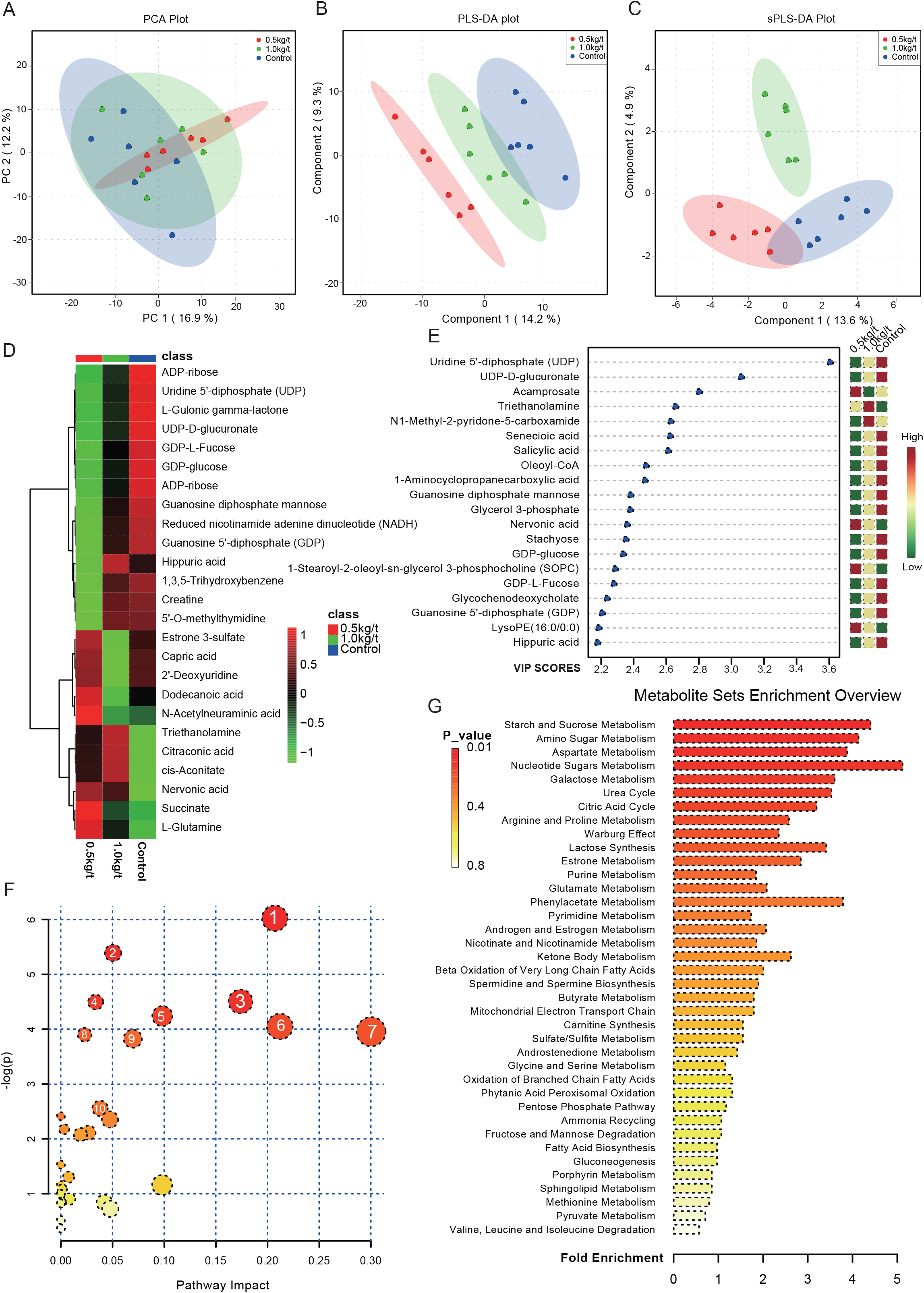
Changes in sow’s breast milk metabolite profile shaped by different lysozyme levels. **A.** Principal components analysis (PCA) plot of metabolic profiles of sow’s milk upon lysozyme supplementation. **B-C.** Partial Least Squares - Discriminant Analysis (PLS-DA) socre plot (**B**) and Sparse Partial Least Squares - Discriminant Analysis (sPLS-DA) plot (**C**)of metabolic profiles and the explained variances are shown in brackets. **D.** Heatmap tree shows metabolite significantly different among groups and their phylogenic relationships. The abundance profiles are expressed by z-scores. **E.** Important features identified by PLS-DA. The colored boxes on the right indicate the relative concentrations of the corresponding metabolite in each group under study. **F.** Bubble diagram of significantly differed pathways. The numbers are in accord with **Table 4** and representing filtered pathways. **G.** Bar Plot for metabolite sets enrichiment. The bar length represent fold enrichiment and the color legend indicating p value.

**Table 3.** Variations in metabolites driven by lysozyme treatment after 21-day supplementation

| Metabolite | VIP | P value<sup>1</sup> | m/z | RT (s) |
| --- | --- | --- | --- | --- |
| L-Glutamine | 1.57525 | 0.004859 | 129.0654 | 93.046 |
| Creatine | 2.59744 | 0.015554 | 132.0767 | 363.362 |
| Triethanolamine | 1.20796 | 0.008983 | 150.1116 | 163.715 |
| 1, 3, 5-Trihydroxybenzene | 1.5075 | 0.016108 | 168.0651 | 430.955 |
| L-Arginine | 2.26854 | 0.029494 | 175.1185 | 523.8465 |
| N-Acetyl-D-glucosamine | 6.81211 | 0.044037 | 204.0863 | 417.33 |
| Argininosuccinic acid | 3.1334 | 0.035875 | 291.1294 | 457.334 |
| S-Methyl-5'-thioadenosine | 3.56887 | 0.046785 | 298.0957 | 67.02 |
| 2'-O-methylcytidine | 1.66055 | 0.038591 | 324.0581 | 436.676 |
| D-Lactose | 31.52 | 0.038208 | 343.1225 | 399.4185 |
| Guanosine 5'-diphosphate (GDP) | 7.40217 | 0.029233 | 444.03 | 439.483 |
| ADP-ribose | 2.9509 | 0.00208 | 560.0759 | 408.2855 |
| GDP-L-Fucose | 4.36646 | 0.026333 | 590.0863 | 439.1075 |
| Reduced nicotinamide adenine dinucleotide (NADH) | 1.03812 | 0.017532 | 666.1275 | 390.424 |
| Stachyose | 1.7266 | 0.02151 | 689.2069 | 457.576 |
| Benzoic acid | 1.38217 | 0.02887602 | 121.0301 | 106.4585 |
| Citraconic acid | 1.77585 | 0.01178885 | 129.0199 | 423.3335 |
| Capric acid | 1.31793 | 0.00196452 | 171.1386 | 47.718 |
| cis-Aconitate | 2.4721 | 0.01471209 | 173.009 | 423.336 |
| Maleic acid | 1.7544 | 0.0143732 | 175.0245 | 352.1695 |
| L-Gulonic gamma-lactone | 1.58526 | 0.001417626 | 177.0399 | 144.86 |
| 2'-Deoxyuridine | 1.60415 | 0.02979523 | 227.0662 | 120.194 |
| Succinate | 2.91345 | 0.003600293 | 235.0448 | 352.1695 |
| D-Ribose 5-phosphate | 1.19978 | 0.04074732 | 289.0314 | 453.871 |
| N-Acetylneuraminic acid | 1.00284 | 0.01040533 | 290.0861 | 341.944 |
| Sucrose | 1.26721 | 0.02556989 | 323.0962 | 429.7805 |
| Estrone 3-sulfate | 1.99979 | 0.03466092 | 349.1089 | 24.8065 |
| Nervonic acid | 1.14023 | 0.01313373 | 365.3397 | 40.47 |
| Uridine 5'-diphosphate (UDP) | 2.21374 | 0.01324003 | 402.9919 | 459.15 |
| Uridine 5'-triphosphate (UTP) | 1.95969 | 0.03726824 | 482.9569 | 430.288 |
| ADP-ribose | 1.48413 | 0.003672366 | 558.0596 | 408.6775 |
| UDP-D-Glucose | 18.7714 | 0.04512712 | 565.0448 | 429.7805 |
| UDP-D-glucuronate | 7.01253 | 0.03221274 | 579.0232 | 459.51 |
| GDP-L-Fucose | 4.89825 | 0.01274555 | 588.0709 | 439.5195 |

1. P values are from one-way ANOVA test.

To explore the biological functions of these metabolic makers, metabolite-set enrichment analysis was performed via MetaboAnalyst v4.0 (Figure 5G). Pathway topology analysis using relative centrality revealed nine significantly different (P < 0.05) enriched pathways (Figure 5F), including alanine, aspartate, and glutamate metabolism, pyrimidine metabolism, arginine and proline metabolism, galactose metabolism, ascorbate and aldarate metabolism, amino sugar and nucleotide sugar metabolism, starch and sucrose metabolism, purine metabolism, and the citrate cycle (Table 4).

**Table 4.** Most enriched pathways of metabolic makers (TOP 10)

| NO. | Pathways | Total | Expected | Hits | P value<sup>1</sup> | Impact |
| --- | --- | --- | --- | --- | --- | --- |
| 1 | Alanine, aspartate and glutamate metabolism | 24 | 0.279 | 3 | 0.002 | 0.207 |
| 2 | Pyrimidine metabolism | 60 | 0.698 | 4 | 0.005 | 0.050 |
| 3 | Arginine and proline metabolism | 77 | 0.896 | 4 | 0.011 | 0.174 |
| 4 | Galactose metabolism | 41 | 0.477 | 3 | 0.011 | 0.034 |
| 5 | Ascorbate and aldarate metabolism | 45 | 0.523 | 3 | 0.014 | 0.098 |
| 6 | Amino sugar and nucleotide sugar metabolism | 88 | 1.024 | 4 | 0.017 | 0.212 |
| 7 | Starch and sucrose metabolism | 50 | 0.582 | 3 | 0.019 | 0.300 |
| 8 | Purine metabolism | 92 | 1.070 | 4 | 0.020 | 0.023 |
| 9 | Citrate cycle (TCA cycle) | 20 | 0.233 | 2 | 0.022 | 0.070 |
| 10 | Butanoate metabolism | 40 | 0.465 | 2 | 0.078 | 0.38 |

1. P values are from Hypergeometric Test.

### Associations between sow’s milk metabolic indicators and gut microbiota mediated by lysozyme supplementation

To further investigate the correlation between breast milk metabolic indicators and the altered gut microbiome driven by lysozyme treatment, Spearman’s correlation coefficients between significantly different metabolites and major genera were also calculated and visualized with heat map (Figure 6A). For instance, L-Glutamine showed significant positive correlations with *Lactobacillus*, *Ruminococcus* 1 and *Lachnospiraceae MK4A136* group. L-Arginine showed significant negative correlations with *Sphaerochaeta*, *Prevotella* 1, *Pseudobutyrivibrio*, and *Prevotella 9*. Moreover, lysozyme-supplementation mediated associations among breast milk composition, serum immunity, and gut microbiota were also summarized and bacteria showed significant impact on both serum biomarkers and breast milk metabolic makers were filtered (Figure 6B).

**Figure.6.**
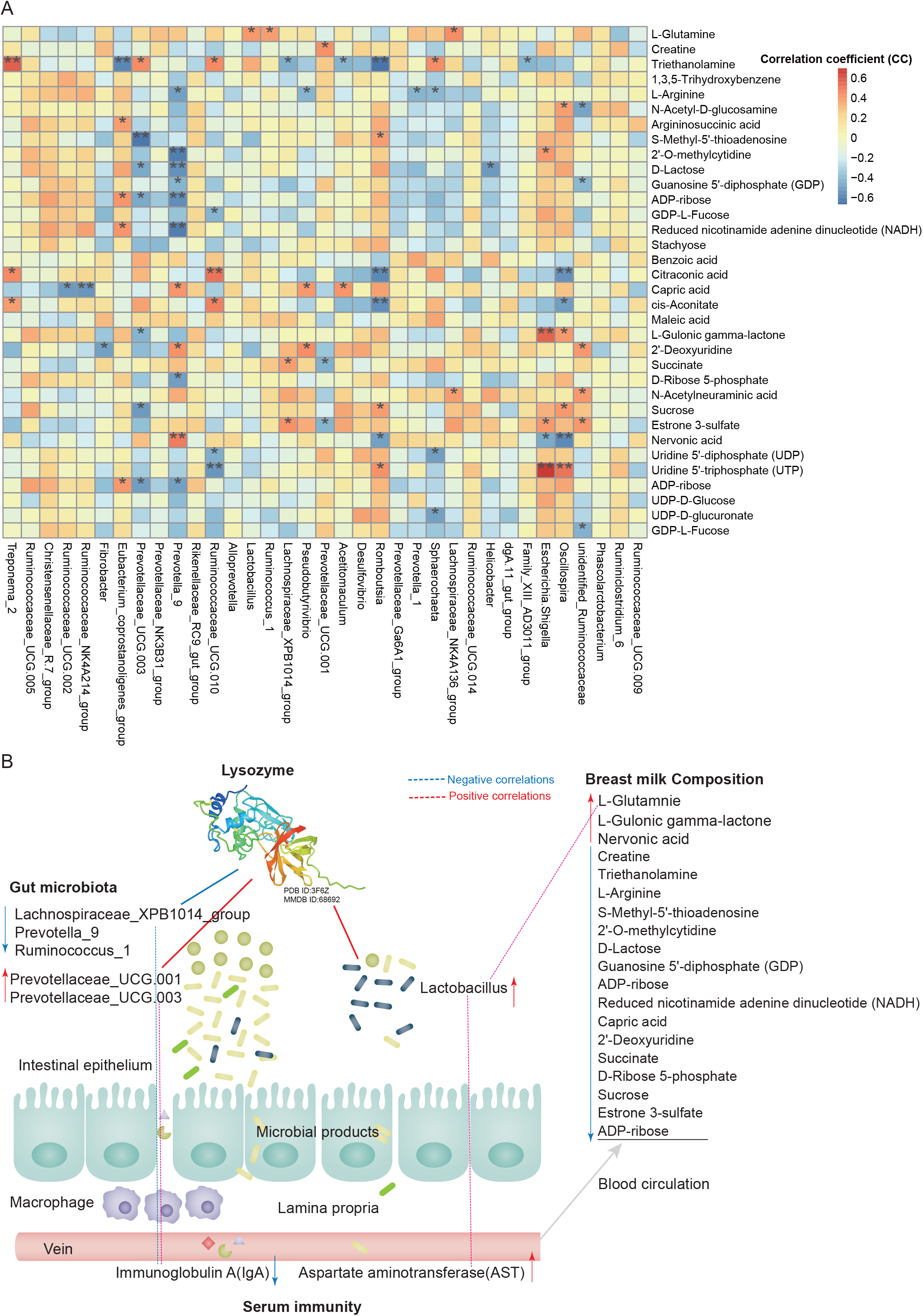
Lysozyme supplementation mediated associations among breast milk composition, serum immunity and gut microbiota. **A.** Correlations between sow’s milk metabiolic indicators and gut microbiota mediated by lysozyme supplementation. **B.** Overview of lysozyme mediated interactions among breast milk composition, serum immunity and gut microbiota revealed by this research.

### Discussion

Recent studies revealed that maternal diets and gut microbiota could directly affect the offspring development early in life [32, 33, 45]. For instance, sows treated with probiotic combinations resulted in improved microbiota diversity in neonatal piglets [46]. In the current study, microbial diversity evidenced by the Shannon index showed a significant reduction in the 1.0 kg/t lysozyme-treated group (p = 0.0014, Figure 1A). No differences in microbial richness were found. Previous studies reported that lysozyme could be effective against a wide range of gastrointestinal pathogens, such as *Listeria monocytogenes*, *Clostridium perfringens*, *Candida* spp., and *Helicobacter pylori* in vitro [7, 21]. Reduced microbial diversity should result from combined factors including physiological preparation for parturition and lysozyme treatment [32, 46]. Furthermore, *Spirochaetes*, *Euryarchaeota,* and *Actinobacteria* significantly increased while *Firmicutes* showed a remarkable reduction in the 1.0 kg/t treated group compared with the control group. *Proteobacteria* also exhibited lower richness in the 1.0 kg/t group (p = 0.077, Table 1). Variations observed in microbial composition driven by lysozymes are in accordance with previous reports. This highlights the effect of lysozymes on beneficial microbe enrichment versus detrimental microbe reduction in the gut microbiome community [2, 11, 47]. In the current research, *Escherichia coli* dramatically reduced in both the 0.5 kg/t and 1.0 kg/t lysozyme-treated groups. ETEC is the major causative agent of diarrhea in weaned pigs, which attaches to the intestinal mucosa, leading to compromised barrier function and malabsorption of large molecules [20, 26, 47]. Previous research also revealed that higher concentrations of lysozyme in milk confer enteric health benefits and prevents ETEC infections in young animals [26, 47, 48, 49]. This study also identified that *Lactobacillus amylovorus* is significantly increased in the 0.5 kg/t group (Figure 1F). *Lactobacillus* is reported to improve the intestinal environment and activate intestinal mucosal immunity in many species, resulting in enhanced SIgA production of the innate immune system, which is essential for the prevention of fimbriae-mediated colonization and the maintenance of intestinal barrier function [3, 50].

Lysozyme driven metabolic function in microbial communities were investigated in this research. The results show that pyrimidine metabolism, purine metabolism, and amino acid related enzymes were significantly upregulated in the 1.0 kg/t lysozyme-treated group. Broadly speaking, lysozyme promoted the shift to greater amino acid and nucleotide metabolism in microbes (Figure 2). In this research, the richness of gram-positive bacteria was significantly down-regulated by lysozyme treatments. Our findings confirmed that lysozyme has a robust antimicrobial activity against gram-positive bacteria, and to a much lesser degree, against gram-negative bacteria [18]. Lysozyme supplementation also significantly increased biofilm formation in the lysozyme-treated groups (Figure 3H). The formation of microbial community biofilms is closely linked to drug resistance and pathogenesis [51, 52]. Further studies may focus on the mechanisms of lysozyme mediated microbial metabolism. In summary, lysozyme supplementation could effectively improve the composition, metabolic functions, and phenotypes of sow’s gut microbiota.

Lysozyme is also an important modulator in non-specific immunity and plays a crucial role in preventing intestinal inflammation [5, 10, 22]. SIgA is predominantly recognized for its role in host defense of the mucosa, where it prevents the invasion of pathogens by neutralization [53]. Lower serum IgA indicates a lower risk of allergy in the postnatal period, which differs from mucosal SIgA [54]. IgM is an important immunoglobulin that functions in the anti-inflammatory response and increased IgM indicating a better immune status [55]. In this study, sow’s serum IgA levels were significantly reduced by increased lysozyme levels. Meanwhile, serum IgM was significantly higher in the 1.0 g/kg group compared with the control group. What is more, serum aspartate transaminase (AST) levels were also significantly upregulated by lysozyme treatment. AST, a liver-specific enzyme that is released into serum following acute liver injury, is used in experimental organ preservation studies as a measure of liver ischemia-reperfusion injury [56]. Hence, reduced serum AST levels induced by lysozyme treatments indicate improved overall health. Lysozyme supplementation could benefit sows with better immune status via up-regulating serum AST and IgM levels.

To determine the effect of dietary lysozyme on sows’ breast milk metabolites, untargeted LC-MS/MS were used to explore the metabolome of all groups. Results showed that metabolites including L-glutamine, succinate, triethanolamine, and L-arginine were significantly upregulated by lysozyme supplementation. It should be noted that L-glutamine could enhance tight junction integrity and proliferation of intestinal porcine epithelial cells [57, 58, 59], which is essential for proper intestinal development. Also, succinate is metabolite that improves glycemic control through the activation of intestinal gluconeogenesis [60]. Our previous work revealed that ethanolamine can enhance intestinal function by altering gut microbiome and mucosal anti-stress capacity [34, 61]. Moreover, L-arginine was shown to play a significant role in shaping the gut microbiota. The intestinal innate immunity of piglets thus improves gut development and protects against pathogenic infection [62, 63, 64]. To sum up, dietary lysozyme supplementation significantly upregulated the beneficial metabolites in sow’s milk, which may exert a long-term impact on the development of offspring. Further, our findings of pathway enrichment and topology analysis of these metabolites revealed nine significantly different (P < 0.05) enriched pathways (Figure 5F) including alanine, aspartate, and glutamate metabolism and pyrimidine metabolism, arginine, and proline metabolism, which further supports the conclusions noted above.

Gut microbiota play multiple roles in animal growth and health, including energy extraction from food, gut barrier function and immune system maturation, and growth performance [30, 31]. For serum immunity makers, 12 genera including *Prevotella*, *Ruminococcaceae* UGG, and *Bacteroides* showed significant correlations with IgA (Figure 4D). *Ruminiclostridium* 9 showed a significant positive relationship with IgM. *Methanobrevibacter* showed a significant negative relationship with AST, while *Lactobacillus* showed significantly positive correlation (Figure 4D). Breast milk metabolites, such as L-glutamine, showed a significant positive correlation with *Lactobacillus*, *Ruminococcus* 1, *Lachnospiraceae*, and the MK4A136 group; and L-arginine showed significant negative correlations with *Sphaerochaeta*, *Prevotella* 1, *Pseudobutyrivibrio* and *Prevotella* 9. These findings confirmed that gut microbiota plays an important role in lysozyme-mediated changes in serum immunity and breast milk composition. Genera that exerted a significant effect on both serum immune indices and milk metabolites were filtered, and a possible regulation network is shown in Figure 6B. Based previous reports, *Lactobacillus* and *Prevotella* may play a key role in lysozyme mediated host-microbial interactions [3, 46, 50].

The application of antibiotics in feeds at subtherapeutic levels could improve performance and overall health and is used extensively throughout the swine industry [15]. Because of the rising concerns about antibiotic resistance and human health, many countries including China (starting in 2020), America, and the European Union banned the use of antibiotics in swine production [1, 15, 24]. Thus, research into alternatives is important for the future development of the livestock industry. As shown in the current study, lysozyme supplementation could effectively improve the composition, metabolic functions, and phenotypes of sow’s gut microbiota and it also benefits sows with better immune status and breast milk composition. Therefore, lysozyme is a viable alternative to traditional subtherapeutic antibiotic use in swine production. Further work will focus on the long-term effect of dietary lysozyme supplementation on commercial swine production and the mechanism of lysozyme mediated host-microbial interactions in cellular models.

## Author contributions

ZJ, XX and Y L designed the study, ZJ, XX, YJ, WK, ZL and SY carried out the animal trials and sample analysis, ZJ, YJ and XX wrote and revised the manuscript.

## Acknowledgements

The authors thank Shanghai E.K.M Biotechnology Company for material assistance.

## Supplemental materials

File named “Supplemental materials”. The assembled HiSeq sequences obtained in the present study were submitted to the NCBI’s Sequence Read Archive (SRA, No. PRJNA415259) for open access.

